# Collaboration of multiple pathways in making a compound leaf

**DOI:** 10.1101/2020.08.19.257550

**Authors:** Alon Israeli, Ori Ben-Herzel, Yogev Burko, Ido Shwartz, Hadas Ben-Gera, Smadar Harpaz-Saad, Maya Bar, Idan Efroni, Naomi Ori

## Abstract

- The variability in leaf form in nature is immense. Leaf patterning occurs by differential growth that occurs during a limited window of morphogenetic activity at the leaf marginal meristem. While many regulators have been implicated in the designation of the morphogenetic window and in leaf patterning, how these effectors interact to generate a particular form is still not well understood.
- We addressed the interaction among different effectors of tomato compound leaf development, using genetic and molecular analyses.
- Mutations in the tomato auxin response factor SlARF5/SlMP, which promotes leaflet formation, suppressed the increased leaf complexity of mutants with extended morphogenetic window. Impaired activity of the NAC/CUC transcription factor GOBLET (GOB), which specifies leaflet boundaries, also reduced leaf complexity in these backgrounds. Analysis of genetic interactions showed that the patterning factors SlMP, GO*B* and the MYB transcription factor LYRATE (LYR) act in parallel to promote leaflet formation.
- This work places an array of developmental regulators in a morphogenetic context. It reveals how organ-level differentiation rate and local growth are coordinated to sculpture an organ. These concepts and findings are applicable to other plant species and developmental processes that are regulated by patterning and differentiation.

## Introduction

Leaf shape ranges from simple leaves, with a single leaf blade, to compound leaves, in which the leaf is composed of separate blade units termed leaflets (Bar & Ori, 2015; Du *et al*., 2018; Efroni *et al*., 2010). Leaf-primordia margins maintain a transient window of morphogenetic activity (Alvarez *et al*., 2016). The elaboration of compound leaves requires a prolonged morphogenetic window (Hagemann & Gleissberg, 1996), and tuning this window enables a species-dependent variability in leaflet number (Blein *et al*., 2010; Chen *et al*., 2010; Hagemann & Gleissberg, 1996; Ori *et al*., 2007; Zhou *et al*., 2014). In tomato, the TCP transcription factor LANCEOLATE (LA) and the MYB transcription factor CLAUSA (CLAU) promote differentiation, thus restricting the morphogenetic window at the leaf margin (Bar *et al*., 2015, 2016; Dengler, 1984; Jasinski *et al*., 2007; Kang & Sinha, 2010; Maltnan & Jenkins, 1962; Ori *et al*., 2007). In contrast, the MYB transcription factor TRIFOLATE (TF) and the tomato KNOTTED1-LIKE HOMEOBOX (KNOXI/TKN2) proteins delay leaf differentiation and preserve the meristematic identity of the leaf margin (Bharathan *et al*., 2002; Blein *et al*., 2013; Chen *et al*., 2010; Hake *et al*., 2004; Hay & Tsiantis, 2009; He *et al*., 2020; Janssen *et al*., 1998; Kimura *et al*., 2008; Naz *et al*., 2013; Peng *et al*., 2011; Shani *et al*., 2009; Zhou *et al*., 2014).

Both simple and compound leaf shape is patterned at the meristematic leaf margin by localized differential growth, producing serrations, lobes, or leaflets (Barkoulas *et al*., 2008; Bar & Ori, 2015; Bilsborough *et al*., 2011; Efroni *et al*., 2010; Nikovics *et al*., 2006; Kawamura *et al*., 2010). The formation of distinct leaflets involves the definition of regions of leaflet initiation and blade growth, alongside intercalary and boundary domains in which growth is inhibited (Fig1a and (Ben-Gera *et al*., 2012, 2016; Bilsborough *et al*., 2011; Blein *et al*., 2008; Fleming, 2006; Koenig *et al*., 2009; Vlad *et al*., 2014)). The plant hormone auxin plays a central role in this patterning mechanism, together with transcription factors such as the CUP-SHAPED COTYLEDONS (CUC) family, which interact with auxin in the domain specification process (Ben-Gera *et al*., 2012; Bilsborough *et al*., 2011; Blein *et al*., 2008; Kierzkowski *et al*., 2019; Xiong *et al*., 2019). In tomato, the Auxin Response Factor (ARF) SlMP/SlARF5 was recently shown acts downstream to auxin to promote leaflet initiation and growth together with additional class A ARF proteins. The activity of these ARFs is antagonized in the intercalary domain by the AUX/IAA protein ENTIRE (E)/IAA9 (Israeli *et al*., 2019). Leaflet initiation and growth is also promoted by the MYB transcription factor LYRATE (LYR) (David-Schwartz *et al*., 2009). While we learned a lot in recent years about the individual functions of these factors in leaf development, it is still not clear how leaf patterning regulators such as auxin and SlMP, LYR and CUC, interact with regulators of the transient morphogenetic window of the leaf margin, such as KNOXI, TF, CLAU and LA. Furthermore, it is not clear how the regulators of the growth, intercalary and boundary domains interact to coordinate leaf patterning.

Here, we examined the genetic and molecular interactions between the described effectors, to investigate how patterning and differentiation interact to achieve leaf shape diversity. We show that an extended morphogenetic window and specific patterning events are both required for stable leaflet production. We further show that the different regulators of the growth and boundary domains act via parallel pathways to pattern leaflets. Therefore, a coordinated network has developed to enable flexible leaf elaboration.

## Materials and methods

### Plant material and growth conditions

Tomato seeds (Solanum lycopersicum cv M82) were germinated and grown for three to four weeks in a growth room or a growth chamber in a 16/8 light/dark regime at 25C^0^ under fluorescent light. Seedlings were then transferred to a greenhouse or to an open field with natural day length and 25^0^C/20^0^C day/night temperature. The *La-2, la-6, clau, Gob4d, gob-3, lyr, slmp-1* and *slmp-2* alleles are from a tomato EMS mutagenesis populations, in the M82 background, and have been described before (Bassel *et al*., 2008; Bar *et al*., 2015; Berger *et al*., 2009; Jasinski *et al*., 2008; Menda *et al*., 2004; Ori *et al*., 2007). The *Pts* (LA2532) and *bip* (LA0663) mutants were obtained from the TGRC and backcrossed to M82. The transgenic lines *35S:Kn1, BLS*≫*TKN2, FIL*≫*miR164, FIL*≫*miR319* and the *DR5:VENUS* were generated in the M82 background and described before (Berger *et al*., 2009; Hareven *et al*., 1996; Ori *et al*., 2007; Shani *et al*., 2009, 2010). EMS Mutant screens were carried out on M82, *La-2/*+ *gob/*+ and *la-6* seeds as described before (Menda *et al*., 2004; Ori *et al*., 2007).

### Trans-activation system

We used the Promoter:LhG4 (p) and Operator (OP) system as described before (Moore *et al*., 1998; Shani *et al*., 2009). Briefly, in this system, driver lines expressing the synthetic transcription factor LhG4 under the control of a specific promoter are crossed to responder lines containing a gene of interest under the control of the E.coli operator, which is recognized by the LhG4 transcription factor but not by any endogenous plant transcription factor. A cross between a driver and a responder line produces an F1 plant in which the selected gene is expressed under the control of the selected promoter (p≫GENE).

### RNA extraction and qRT-PCR analysis

RNA was extracted using the Plant/Fungi Total RNA Purification Kit (Norgen Biotek, Thorold, ON, Canada) according to the manufacturer’s instructions, including DNase treatment. cDNA synthesis was performed using the Verso cDNA Kit (Thermo Scientific, Waltham, MA, USA) or SuperScript II reverse transcriptase (18064014; Invitrogen, Waltham, MA, USA) using 1 mg of RNA. qRT-PCR analysis was carried out using a Corbett Rotor-Gene 6000 real-time PCR machine, with SYBR Premix for all other genes. Levels of mRNA were calculated relative to the EXPRESSED (EXP) or TUBULIN (TUB) genes as internal controls as follows: in each biological repeat, the expression levels of the assayed gene and EXP/TUB were separately calculated relative to a standard curve obtained by a dilution series of a reference sample. The gene expression level in each biological repeat was calculated by dividing the gene expression value by that of EXP/TUB. Average expression values were then calculated and presented as ‘relative gene expression’. Each biological repeat included between 3-15 leaf primordia, depending on the developmental stage. Primers used for the qRT-PCR analysis are detailed in Table S2.

### Phenotyping, Imaging and Scanning Electron Microscopy (SEM)

Images analyzing the early developmental stages of the whole leaf primordium were captured using an Olympus SZX7 stereo microscope (http://www.olympus.com/) equipped with a Nikon DXM1200 camera and ACTA software, or a Nikon SMZ1270 stereo micro- scope equipped with a Nikon DS-Ri2 camera and NIS-ELEMENTS software. The expression pattern of the *DR5::VENUS* reporter was detected by a Stereomicroscope, as described before (Shani *et al*., 2010; Bar *et al*., 2016). For scanning electron microscopy (SEM), tissues were fixed in 30% Ethanol and vacuumed for 10 min, followed by dehydration in an increasing ethanol series up to 100% ethanol. Fixed tissues were critical-point dried, mounted on a copper plate and coated with gold. Samples were viewed using a JEOL 5410 LV microscope (Tokyo, Japan).

### Leaf quantification

Phenotyping and quantification of leaf form, petiole length and shoot architecture were performed on field- or greenhouse-grown plants. Collected Representative mature intact leaves or mature plants, were photographed using a Nikon D5200 camera and the photographs used for quantification of leaf-shape phenotype. Leaflet order was defined, and leaflet number quantified as described before (Bar *et al*., 2015; Shani *et al*., 2010; Yanai *et al*., 2011). Briefly, primary leaflets are separated by a rachis, and some of them develop secondary and tertiary leaflets. Intercalary leaflets are lateral leaflet that develop from the rachis later than the primary leaflets and between them. Each genotype was represented by at least 3 biological replicates, consisting of leaves from different plants. Mean values were statistically analyzed using the Student’s t-test. Student’s t-test (two-tailed) was used for comparison of means, which were deemed significantly different at pv - 0.05. Images were manipulated uniformly using adobe Photoshop.

### Accession Numbers

Sequence data used in this study can be found in the Sol Genomic Network under the following accession numbers: SlMP - solyc04g081240; ENTIRE/SlIAA9 - Solyc04g076850; LYR - Solyc05g009380; GOBLET - Solyc07g062840; LANCEOLATE - Solyc07g062680; CLAUSA - Solyc04g008480; TKN2 - Solyc02g081120; TRIFOLIATE - Solyc05g007870; PETROSELINUM - Solyc06g072480.

## Results

### SlMP and LYRATE promotes growth in parallel pathways

We have previously shown that the tomato ARF transcription factor SlMP/SlARF5 promotes organ initiation and growth (Israeli *et al*., 2019). Previous studies have shown that the MYB transcription factor LYRATE (LYR) also promotes leaflet initiation and blade growth (Bar *et al*., 2015; David-Schwartz *et al*., 2009). *lyr* mutants have less primary and intercalary leaflets, similar to *slmp* mutants (Fig 1), while plants that overexpress *LYR* have ectopic blade growth in the intercalary domain, similar to *entire* (*e*) mutants, in which auxin response and SlMP/SlARF5 activity are enhanced in the intercalary domain (David-Schwartz *et al*., 2009). We examined how SlMP/SlARF5 and LYR interact to promote the growth domain, and whether they act in the same genetic pathway, by investigating their genetic and molecular interactions. Strikingly, *lyr* and *slmp* enhanced each other, with the double mutants showing further reduction in primary leaflet number in comparison to the single mutants (Fig 1b-e). *lyr slmp* leaves had a range of phenotypes, with the most severe leaves being flattened, nearly bladeless, and lacking leaflets (Fig 1b-e and S1). The most severe phenotypes occurred in early leaves and leaves of axillary branches. The substantial enhancement of the single mutants suggests that SlMP and LYR promote leaflet initiation and growth via parallel pathways (Fig 1o). It was previously shown that *lyr* partially suppresses the ectopic blade phenotype of *e* in the VF36 background (David-Schwartz *et al*., 2009). We generated a *e lyr* double mutant in the M82 background and compared it to the *e slmp* double mutant. This comparison highlighted the differences between the two double mutants: While *e* and *slmp* mutually suppressed each other, restoring a wild-type leaf form (Israeli *et al*., 2019), *lyr* only partially suppressed the ectopic blade growth of *e*, mainly in the basal region of the leaf (Fig 1f-h). We further investigated the effect of these two mutants on auxin response, by comparing the effect of the single and double mutants on the expression of the auxin response marker *DR5*. While in wild-type leaf primordia *DR5* is expressed specifically in the growth domain, marking leaflet initiation sites (Fig 1i and (Shani *et al*., 2010)), in the *slmp* mutant *DR5* expression is expanded into the intercalary domain (Fig 1j and (Israeli *et al*., 2019)). In contrast, in the *lyr* mutant *DR5* expression was comparable to that of the wild type, and in *lyr slmp* double mutants, *DR5* expression was similar to single *slmp* mutants, despite the reduced blade growth domain (Fig 1k-l). In agreement with the genetic and *DR5* analyses, we did not detect an effect of the *lyr* and *slmp* mutants on the expression of *SlMP* and *LYR*, respectively (Fig 1m, n). Therefore, we concluded that SlMP and LYR promote leaflet initiation and growth via separate pathways and the lack of both severely impairs marginal growth (Fig 1o).

**Fig 1.**
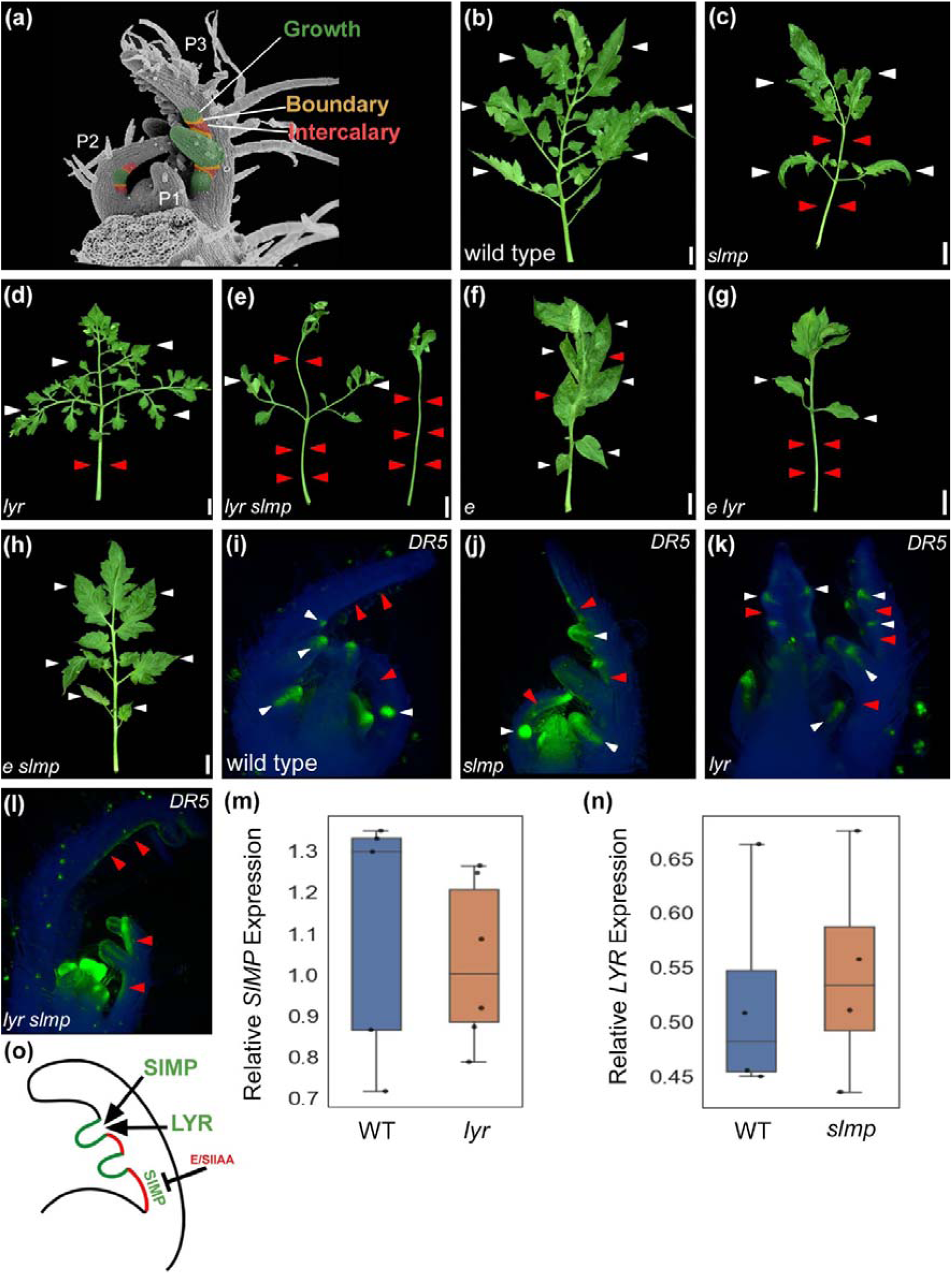
SlMP and LYR promote growth in parallel pathways. (A) Scanning Electron Microscope (SEM) image of a wild-type shoot apex containing the Shoot Apical Meristem (SAM) and the three youngest leaf primordia (m+P3), color-coded for the growth domain (green), boundary domain (orange) and intercalary domain (red). (B-H) 5^th^ leaf of the indicated genotypes. Scale bars: 2 cm. White and red arrowheads represent primary leaflets and missing primary and intercalary leaflets, respectively. (I-L) Stereomicroscope images showing the distribution of the *DR5:VENUS* marker in the SAM and young leaf primordia of the indicated genotypes. (M-N) qRT-PCR analysis of the mRNA expression of the indicated genes, relative to the *EXP* reference gene in 5^th^ leaf primordia of the indicated genotypes. In all the boxplots the central horizontal line marks the median, upper and lower box lines represent the first and third quartile, respectively, and the whiskers represent the maximum and minimum values of at least three biological replicates, each containing between 1–3 plants. (O) A schematic model for the proposed interaction between E, SlMP and LYR in the regulation of the growth and intercalary domains. E inhibits growth in the intercalary domain (red) by repressing the activity of SlMP. SlMP and LYR promote the growth domain (green) via parallel pathways.

### SlMP and GOBLET jointly promote leaflet formation

The tomato CUC gene *GOBLET* (*GOB*) plays an important role in boundary specification at the leaf margin, and altered GOB/CUC2 expression or activity substantially alters leaf patterning (Fig 2, 5 and (Ben-Gera *et al*., 2012; Berger *et al*., 2009)). Plants overexpressing *microRNA164* (*MIR164*), which targets *GOB/CUC2*, or loss-of-function *gob-3* mutants, have less leaflets and smooth leaf margins (Fig 2g, 5c red and purple arrowheads and (Berger *et al*., 2009; Brand *et al*., 2007)). The gain-of-function mutant *Gob4d/*+, which has a point mutation in the *MIR164* target site leading to loss of *MIR164* recognition and thus deregulation, has deeply lobed leaflets (Fig 2e, 5i blue arrowheads and (Berger *et al*., 2009; Brand *et al*., 2007;)). In Arabidopsis, CUC2, auxin and auxin transport regulate serrations in a coordinated feedback loop: CUC2 promotes serrations by regulating auxin transport, and auxin in turn represses CUC2 (Bilsborough *et al*., 2011; Galbiati *et al*., 2013). To understand how these tools are utilized and coordinated in the context of compound-leaf development and leaflet formation, we examined the molecular and genetic interaction between SlMP and GOB/CUC2. We first analyzed the effect of SlMP and GOB activities on the expression of each other. *SlMP* expression was similar between wild type, *Gob4d, FIL*≫*GOB*^*m*^, and *FIL*≫*miR164* (Fig 2a). Similarly, *GOB* expression was not affected by the *slmp* mutant (Fig 2b). *Gob4d* and *slmp* both have elongated petioles and less leaflets (Fig 2c-e and i, j). In agreement with the expression analysis, *slmp Gob4d/*+ double mutants had partially additive phenotypes: *slmp Gob4d/*+ double mutants had fewer primary leaflets and longer petioles than did each of the single mutants (Fig 2c-f and i, j). *slmp FIL*≫*miR164* plants also showed additive phenotypes (Fig 2g, h). These results suggest that SlMP-mediated leaflet initiation and GOB/CUC2-mediated boundary specification are parallel pathways acting together to determine the number and location of leaflets (Fig 2k). Therefore, GOB/CUC2 affects leaflet patterning not only through auxin. The effect of GOB/CUC2 appears to differ between lobe patterning and leaflet patterning. The effect on lobes is conserved with Arabidopsis, while the effect on leaflets is different, suggesting that leaflet and lobe patterning utilizes similar tools in a different manner.

**Fig 2.**
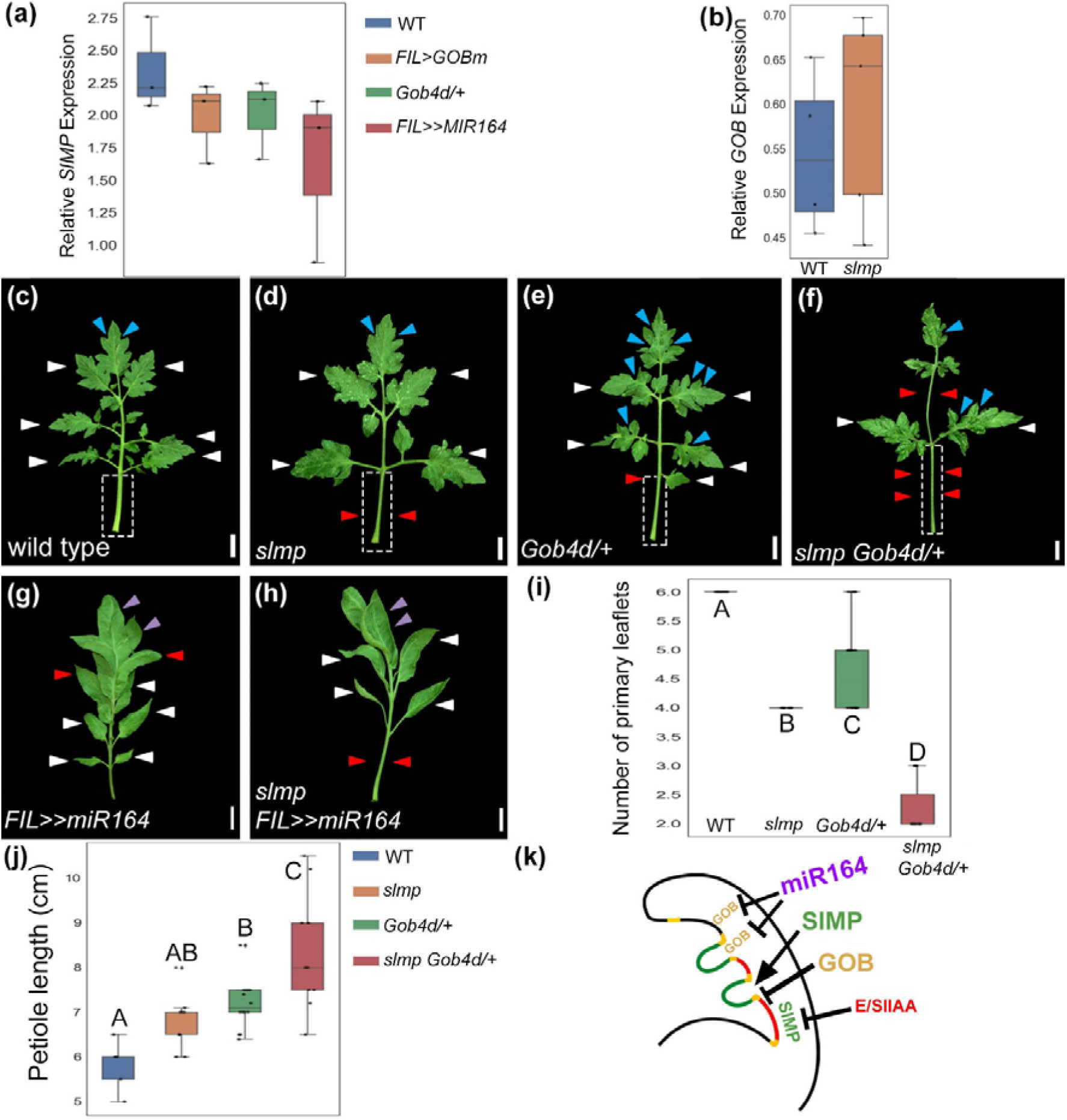
SlMP promotes and GOB restricts growth in parallel pathways. (A-B) qRT-PCR analysis of the mRNA expression of the indicated genes, relative to the *EXP* reference gene in 5^th^ leaf primordia of the indicated genotypes. Boxplots of at least three biological replicates, each containing between 1–3 plants. (C-H) 5^th^ leaves of the indicated genotypes. White, red, blue and purple arrowheads represent primary leaflets, missing leaflets, deep lobes, and smooth leaf margin, respectively. The dashed squares outline the leaf petioles, as quantified in J. Scale bars: 2 cm. (I) Quantification of the number of primary leaflets in the indicated genotypes. Boxplots of 5–10 leaves. Different letters indicate statistical significances between the indicated genotpyes, by Tukey– Kramer multiple comparison statistical test, P < 0.05. (J) Quantification of the petiole length of the indicated genotypes. Boxplots represent the SE of 5–10 leaves. Different letters indicate statistical significances between the indicated genotpyes, by Tukey– Kramer multiple comparison statistical test, P < 0.05. (K) A schematic model for the proposed interaction between SlMP and GOB in the regulation of growth at the leaf margin. SlMP promotes and GOB inhibits growth in the growth and boundary domains, respectively. The temporal and spatial specification of both domains determine leaflet number and location.

### SlMP promotes leaflet formation within a transient developmental window

The results above suggest that several regulators act in parallel within the same domain (MP and LYR), and that different patterning factors (MP and GOB) act in parallel to promote the growth and intercalary domain, respectively. We then asked how these patterning processes are integrated with the morphogenetic activity of the leaf marginal meristem. First, we studied in more detail *SlMP* expression during leaf development (Fig 3). Successive leaves vary in their complexity and maturation rate, such that at the P5 stage, the first leaf (L1) has matured and ceased generating leaflets, while later leaves still make leaflets (Bilsborough *et al*., 2011; Shleizer-Burko *et al*., 2011). *SlMP* expression showed a gradual increase in P5 primordia of successive leaves and peaked earlier in leaf development in the first leaf, in agreement with a role in a specific developmental window of leaflet formation (Fig 3a, b). In addition, we found higher expression of *SlMP* in the margins of P7-P8 compared with inner tissues (Fig 3c, d). We therefore assessed the effect of *slmp* on the number of leaflets in successive leaves. All *slmp* leaves showed a reduction in the number of leaflets compared to the wild type, and the effect increased with the increase in leaf complexity (Fig 3e, f, g). This suggests that *SlMP* acts in the leaf margin at a specific spatial and temporal context to promote leaflet initiation and growth.

**Fig 3.**
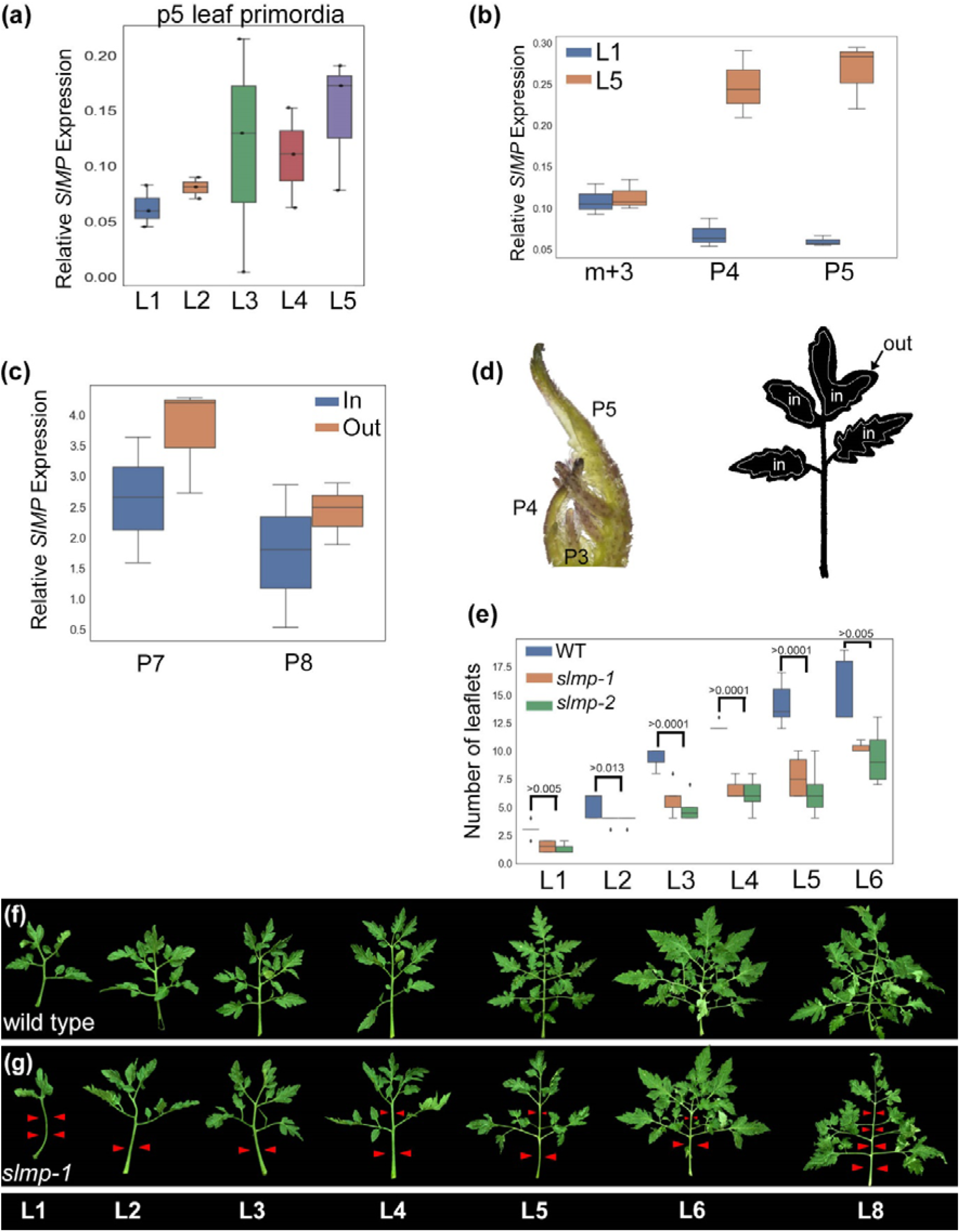
The spatial and temporal expression of *SlMP* correlates with leaflet formation. (A-C) qRT-PCR analysis of the mRNA expression of *SlMP*, relative to the *EXP* reference gene. B shows the expression at the P5 stage in primordia of successive leaves, where L1 is the first leaf produced by the plant. C shows the expression at different developmental stages of the first (L1) and fifth (L5) leaves produced by the plant. m+3 - SAM and the 3 youngest primordia, in which L1 or L5 were at the P3 stage. D shows the expression in the margin (out) and inner parts of the leaf (in), as illustrated in E, of the fifth leaf at the P7 and P8 stages, as indicated. Boxplots of at least three biological replicates, each containing at least 3 plants. (D) Stereomicroscope image showing the meristem and the five youngest leaf primordia of wild-type (left), the tissue used for the quantification of *SlMP* expression in (B-C), and a silhouette (right) showing the dissected domains used for the quantification of *SlMP* expression shown in (D). (E) Effect of two *slmp* alleles on the number of primary and intercalary leaflets in succesive leaves. At least 7 plants were included for each genotype. The P-value for the statistical significance of the difference between the wild type and *slmp*, by Tukey–Kramer multiple comparison statistical test, P < 0.05, is shown for each leaf. (F-G) Successive leaves of wild type (A) and *slmp-1* (B). The leaf number is indicated at the bottom. Red arrowheads point to missing leaflets. Scale bars: 2 cm. The leaves are not to scale.

We genetically examined this idea by studying the relationship between SlMP and regulators of the morphogenetic window. *LA/TCP4* restricts the morphogenetic potential of tomato leaves by promoting leaf differentiation (Nancy G. Dengler, 1984; Ori *et al*., 2007). The dominant, gain-of-function mutant *La-2* shows accelerated leaf maturation and differentiation, resulting in small leaves and reduced leaflet formation (Fig 4a, b and (Ori *et al*., 2007)). Conversely, the MYB transcription factor TRIFOLIATE (TF) promotes morphogenesis and delays differentiation, and leaf development terminates preciously in *tf* loss-of-function mutants leading to small leaves with only one terminal and two lateral leaflets (Fig S2i and (Naz *et al*., 2013)). Leaves of *La-2/*+ *slmp* double mutants were similar to those of *La-2/*+, but were slightly larger with longer petioles, which are characteristics of single *slmp* mutants (Fig 4a-d, S2l). Similarly, *tf slmp* double mutants were almost identical to *tf* single mutant, but slightly larger (Fig S2i, j, l). This finding is in agreement with the expression dynamics of *SlMP* (Fig3) and suggests that *La-2/*+ and *tf* leaves mature fast and cease morphogenesis before the window of SlMP activity. In agreement, *SlMP* expression was reduced in *La-2/*+ primordia compared with the wild type (Fig 4i).

**Fig 4.**
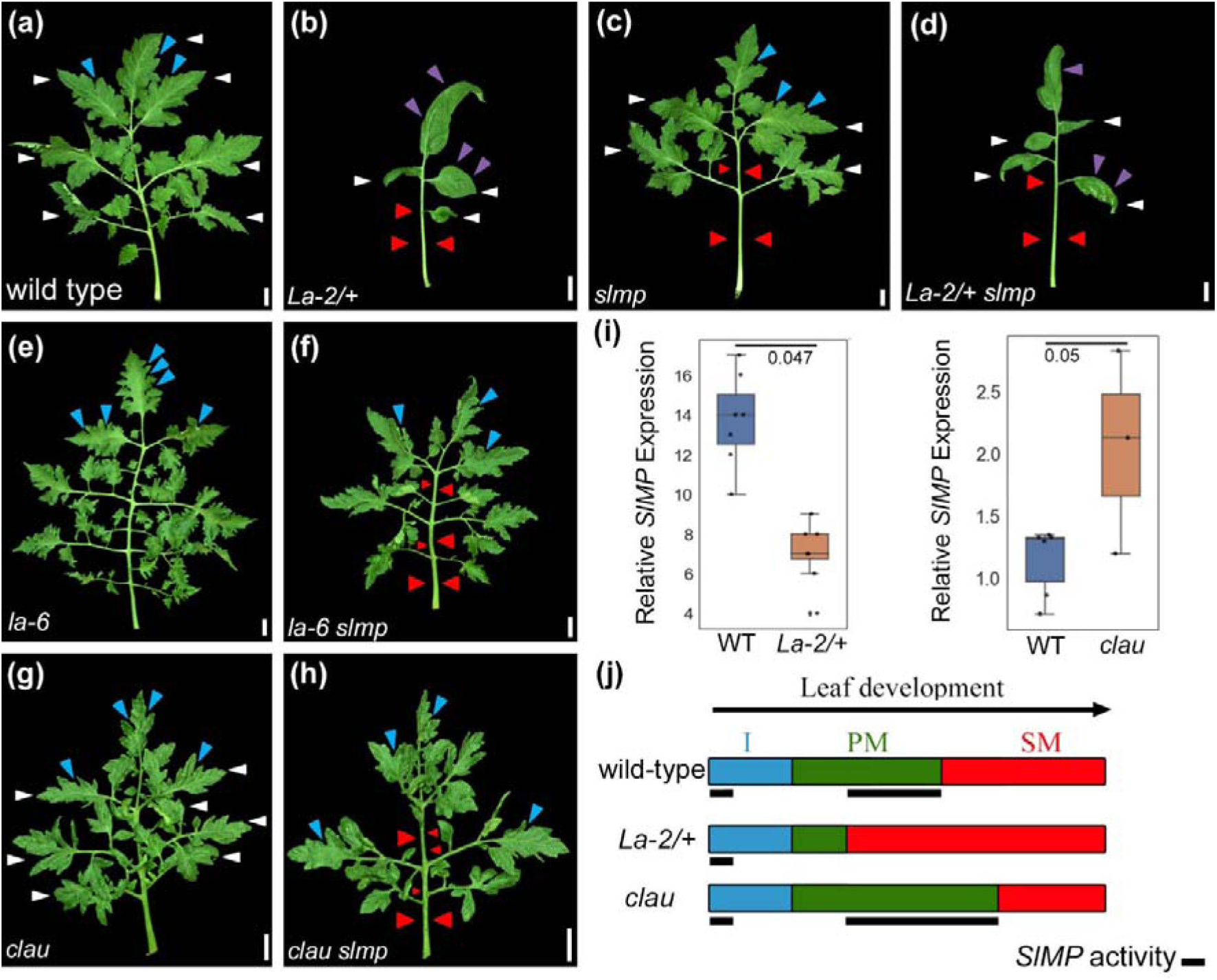
SlMP promotes leaflet formation within a defined morphogenetic window. (A-H) 5^th^ leaf of the indicated genotypes. White, red, blue and purple arrowheads represent primary leaflets, missing primary and intercalary leaflets, dissected lobes and smooth leaf margin, respectively. Scale bars: 2 cm. (I) qRT-PCR analysis of the mRNA expression of *SlMP*, relative to the *EXP* reference gene in the indicated genotypes. Boxplots represent the SE of at least three biological replicates, each containing at least 3 plants. Significant differences by Student’s t test. (J) A schematic illustration describing the progress of leaf development. The color codes of the different stages are indicated in the wild type: Initiation (I) in blue, Primary Morphogenesis (PM) in green and Secondary Morphogenesis (SM) in red. Black lines designate SlMP activity during leaf and leaflet initiation. In the wild type, LA and CLAU promote differentiation. Gain-of-function *La-2/*+ leades to early differentiation, shorter PM, and a consequent reduction in *SlMP* expression, which acts at this stage to promote leaflet formation. By contrast, *clau* mutation results in an extended PM stage and a consequent increase in *SlMP* expression and increased leaflet formation and leaf eomplexity. The increased leaf complexity is mediated in part by SlMP.

To further characterize the spatial and temporal context of SlMP activity, we generated double mutants between *slmp* and several mutants with increased leaf complexity. The loss-of-function mutants *la-6* and *clau* have elaborated and dissected compound leaves due to extended morphogenetic activity (Fig 4e, g and (Avivi *et al*., 2000; Bar *et al*., 2016)). *la-6 slmp* and *clau slmp* double mutants showed reduced complexity relative to the single *la-6* or *clau* mutants (Fig 4e-h and Fig S2m, missing leaflets are marked with red arrowheads). *slmp* also substantially suppressed the increased leaflet formation of mutants and transgenic lines with elevated expression or activity of *KNOXI* genes, including *Me/*+, *BLS*≫*TKN2, bippinata* (*bip*) and *Petroselinum (Pts)* (Fig S2a-h, m and (Kimura *et al*., 2008; Parnis *et al*., 1997; Shani *et al*., 2009)). In agreement, we found elevated expression level of *SlMP* in *clau* leaf primordia (Fig 4i). Interestingly, *TKN2* expression decreased in *slmp* leaf primordia, suggesting a more complex interaction between these factors (Fig S2k). Therefore, mutants that shorten the morphogenetic window are epistatic to *slmp*, while increased leaf complexity of mutants that prolong the morphogenetic window depends on intact SlMP. Cumulatively, these results indicate that marginal activity largely depends on SlMP-mediated leaflet formation and that similar developmental programs mediate initiation and growth from the leaf margins.

### GOBLET specifies leaflet boundaries within a transient developmental window

To genetically examine the respective contribution of boundary specification and maturation rate to leaf shape, we introgressed the different *gob* alleles into genotypes with altered morphogenetic windows. *MIR164* overexpression suppressed the *la-6* increased leaflet number and lobing (Fig 5a-e, red and blue arrowheads). Strikingly, expression of *MIR164* substantially suppressed the phenotype of the super compound leaves and undifferentiated margins caused by overexpression of MIR319, which targets LA/TCP4 and three additional class II TCPs (Fig S3a-c). Similarly, *MIR164* overexpression or *gob-3* suppressed the increased dissection caused by overexpression of the maize KNOXI gene *kn1* or by *Me/*+, respectively (Fig 5f, g and S3d-i). This suggests that proper specification of the boundary domain by GOB/CUC2 is essential for the enhanced complexity that results from a prolonged morphogenetic window (Fig 5 and S3). In contrast, *La-2/*+ was nearly epistatic to alteration of GOB/CUC2 activity in *La-2/*+ *FIL*≫*MIR164* and *La-2/*+ *Gob4d/*+ leaves, although the increased and decreased lobing caused by *Gob4d/*+ and *FIL*≫*MIR164*, respectively, were still apparent in *La-2/*+ *Gob4d/*+ and in *La-2/*+ *FIL*≫ *MIR164* (Fig 5h-k). These results suggest that GOB/CUC2 acts within the morphogenetic window that is defined partially by LA/TCP4 to specify leaflet boundaries. However, the effect of GOB/CUC2 on lobing is less dependent on the morphogenetic window than the number of leaflets, or LA/TCP4 has a more prominent role in leaf maturation than in leaflet maturation. Together, these genetic interactions suggest that marginal activity depends on both a prolonged morphogenetic window at the leaf margin, and proper specification of growth and boundary domains within this window.

**Fig 5.**
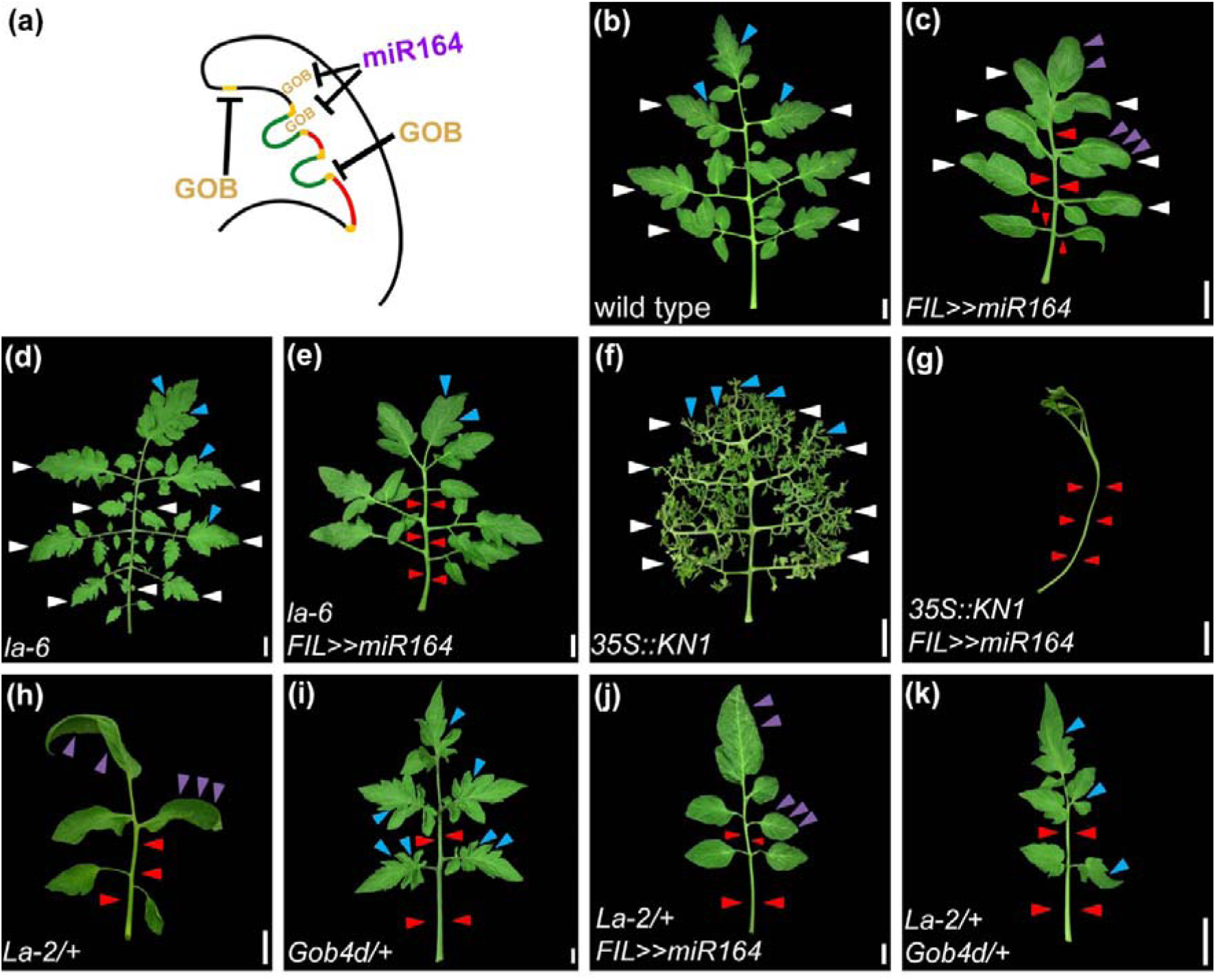
GOBLET defines leaflet boundairs within a defined morphogenetic window. (A) A schematic model for the proposed role of GOBLET in growth inhibition in the boundary domain. (B-K) 5^th^ leaves of the indicated genotypes. Scale bars: 2 cm. White, red, blue and purple arrowheads represent primary leaflets, missing leaflets, deep lobes, and smooth leaf margin, respectively.

### Prolonged morphogenetic window is essential for leaf complexity

The genetic interactions presented above suggest that a prolonged morphogenetic window is essential for the manifestation of leaflet patterning events. To examine this idea in a broader view, we performed genetic screens in two genetic background with opposite effects on the morphogenetic window: The *La-2* mutant, with rapid maturation and a short morphogenetic window and the *la-*6 mutant with prolonged maturation and extended morphogenetic window. We hypothesized that the *La-2* background will be relatively insensitive to the identification of new patterning regulators, while *la-6* will serve as a sensitive background for the potential identification of new modifiers. We previously mutagenized *La-2/*+ *gob/*+ progeny with ethyl methane sulfonate (EMS), a population that we used for the identification of the *la-6* loss-of-function allele (Menda *et al*., 2004; Ori *et al*., 2007). We have screened ∼1800 M2 families from this population, however, very few mutants that affected the *La-2* leaf phenotype were identified (Fig 6 and S4), in agreement with *La-2/*+ being a relatively insensitive background for the identification of leaf shape mutants. Three *La-2/*+ enhancers (h1413, h586 and h1241 the latter two being alleles of the same gene) were identified (Fig 6, S4 and table 1). In contrast to the screen in the *La-2/*+ background, from the ∼1800 M2 families that were screened in the *la-6* background, we identified 37 suppressors and 14 enhancers of the *la-6* leaf phenotype (Fig 6, S5 and table 1). Therefore, the *La-2/*+ genetic background appears epistatic to mutations in many potential patterning regulators (similarly to *slmp* and *gob-3*) and the *la-6* genetic background enabled the identification of potential new patterning regulators. In general, this forward genetics, unbiased approach strongly supports the importance of both extended morphogenesis and marginal patterning for leaf-shape diversity.

**Fig 6.**
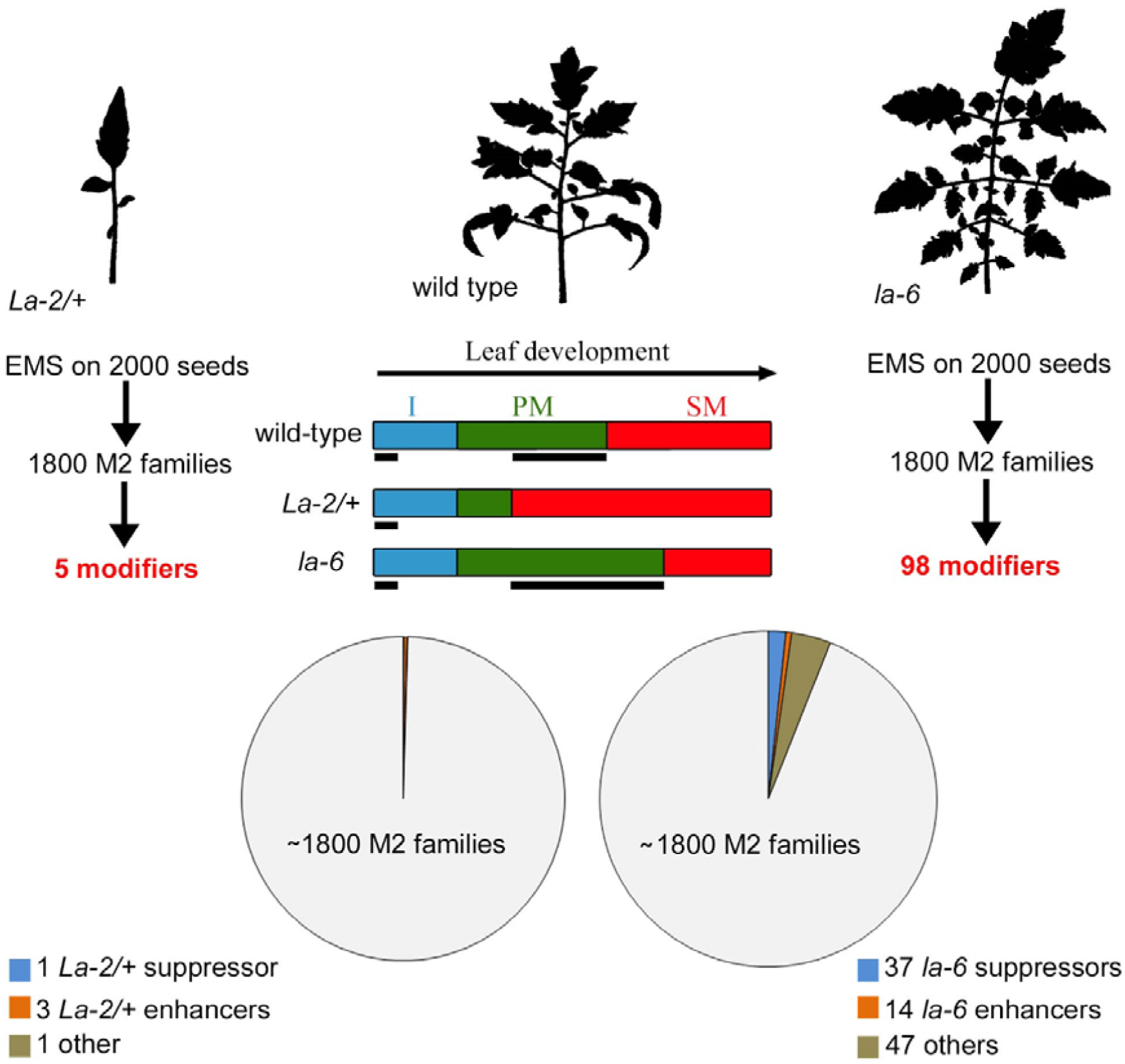
Extended morphogenetic window allows the manifestation of genetic modifiers. Top: Silhouettes of representative 5^th^ leaves of the indicated genotypes. *La-2/*+ and *la-6* were used as backgrounds for 2 independent mutant screens. Below the wild type leaf silhouette is a scheme showing the relative length of the developmental windows in these genotypes. Below each mutant is a scheme describing the mutant screen and the number of enhancers and suppressors identified.

**Table 1.** mutants from the *la-6* suppressors screen that were identified during this study.

| Plant number | Phenotype |
| --- | --- |
| 104 | la suppressor |
| 284 | la suppressor |
| 287 | la suppressor |
| 387 | la suppressor |
| 389 | la suppressor |
| 425 | la suppressor, simple leaf |
| 453 | large leaf |
| 472 | la suppressor |
| 487 | strong la suppressor |
| 510 | la suppressor |
| 517 | weak la suppressor |
| 529 | la suppressor |
| 586 | la suppressor |
| 613 | la suppressor |
| 619 | la suppressor |
| 641 | la suppressor |
| 664 | la suppressor |
| 682 | la suppressor |
| 738 | la suppressor |
| 749 | dwarf, flowers on leaf |
| 756 | la suppressor |
| 955 | dwarf |
| 956 | la suppressor |
| 977 | la suppressor |
| 979 | la suppressor |
| 997 | la suppressor |
| 1221 | la suppressor |
| 1222 | dwarf |
| 1250 | large blade |
| 1346 | la suppressor |
| 1352 | la suppressor |
| 207 | la enhancer |
| 611 | la enhancer |
| 651 | la enhancer |
| 681 | la enhancer |
| 685 | lateral suppressor, la enhancer, lobed cotyledons |
| 708 | late flowering, la enhancer, chlorotic |
| 1271 | growth arrest |
| 17 | sp enhancer |
| 101 | sp enhancer |
| 359 | sp enhancer |
| 360 | sp enhancer, simple leaf, la suppressor |
| 509 | sp enhancer |
| 836 | sp enhancer |
| 1185 | sp enhancer, simple leaf |
| 191 | la suppressor |
| 585 | sp suppressor |
| 1175 | sp suppressor |
| 72 | anthocyanin accumulation, la suppressor |
| 73 | light green, divided meristem |
| 1054 | chlorotic |
| 1263 | chlorotic |
| 1361 | chlorotic, irregular growth |
| 44 | dwarf, less leaflets, simple leaf |
| 107 | dwarf |
| 302 | supper dwarf |
| 735 | dwarf |
| 951 | dwarf, deep lobes |
| 1149 | dwarf |
| 1320 | supper dwarf, lobed cotyledons |
| 194 | lateral suppressor |
| 265 | weak lateral suppressor |
| 346 | narrow leaf |
| 494 | narrow leaf |
| 506 | narrow leaf, many leaflets |
| 372 | large leaf, round leaflets |
| 507 | large leaf |
| 573 | large leaf |
| 596 | long petiole |
| 638 | large leaf |
| 963 | large leaf |
| 1041 | large leaf |
| 1063 | large blade |
| 1158 | large blade |
| 1253 | large leaflets |
| 1270 | large leaf, yellow strips |
| 1273 | large leaf |
| 1396 | large leaf |
| 205 | deep lobes |
| 635 | lobed cotyledon |
| 1224 | less dissected, lobed |
| 1362 | lobed cotyledons |
| 355 | growth arrest |
| 974 | growth arrest, early flowering |
| 1207 | growth arrest |
| 525 | growth arrest |
| 971 | growth arrest |
| 1285 | la suppressor |
| 430 | wirey |
| 1101 | wirey |
| 338 | strange leaves |
| 713 | round leaflets |
| 841 | trifoliate like |
| 940 | Lanceolate like |
| 1071 | folded leaf |
| 1256 | simple round leaflets |
| 1303 | strange leaves |
| 584 | slmp-1 |

**Table 2.**
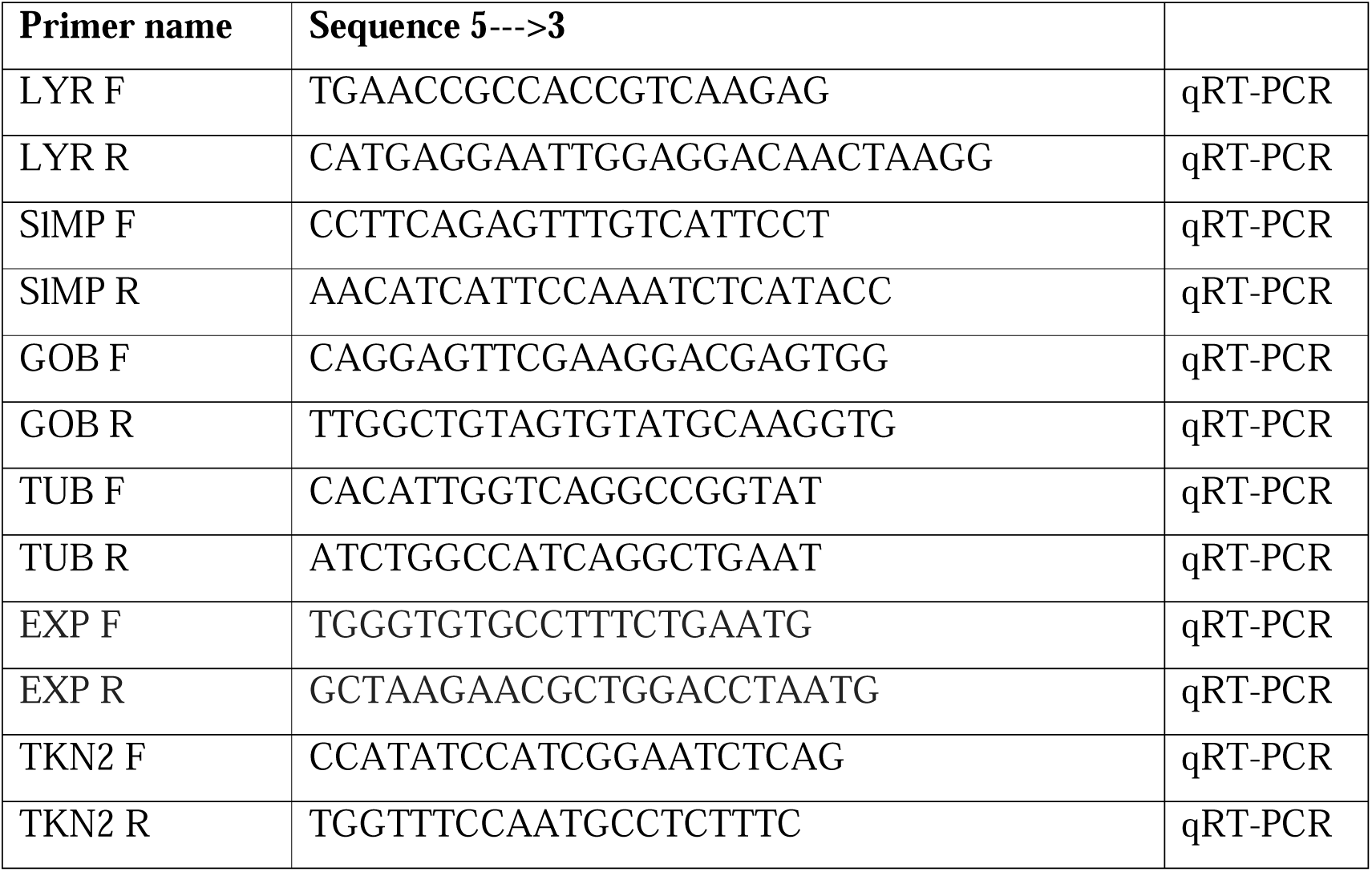
qRT-PCR primers used in this study.

## Discussion

Morphological diversity is achieved by the combination of tuning global differentiation in time and space, and patterning by local growth (Kierzkowski *et al*., 2019; Tsukaya, 2019). Here, using the tomato compound leaf as a model, we examined how these two processes cooperate to pattern the leaf and generate leaf-shape diversity. We found that coordinated specification of growth and boundary domains takes place within a transient morphogenetic window, jointly enabling the elaboration of leaflets and lobes (Fig 7). Within each domain, several pathways act in parallel to ensure robust shape patterning.

**Fig 7.**
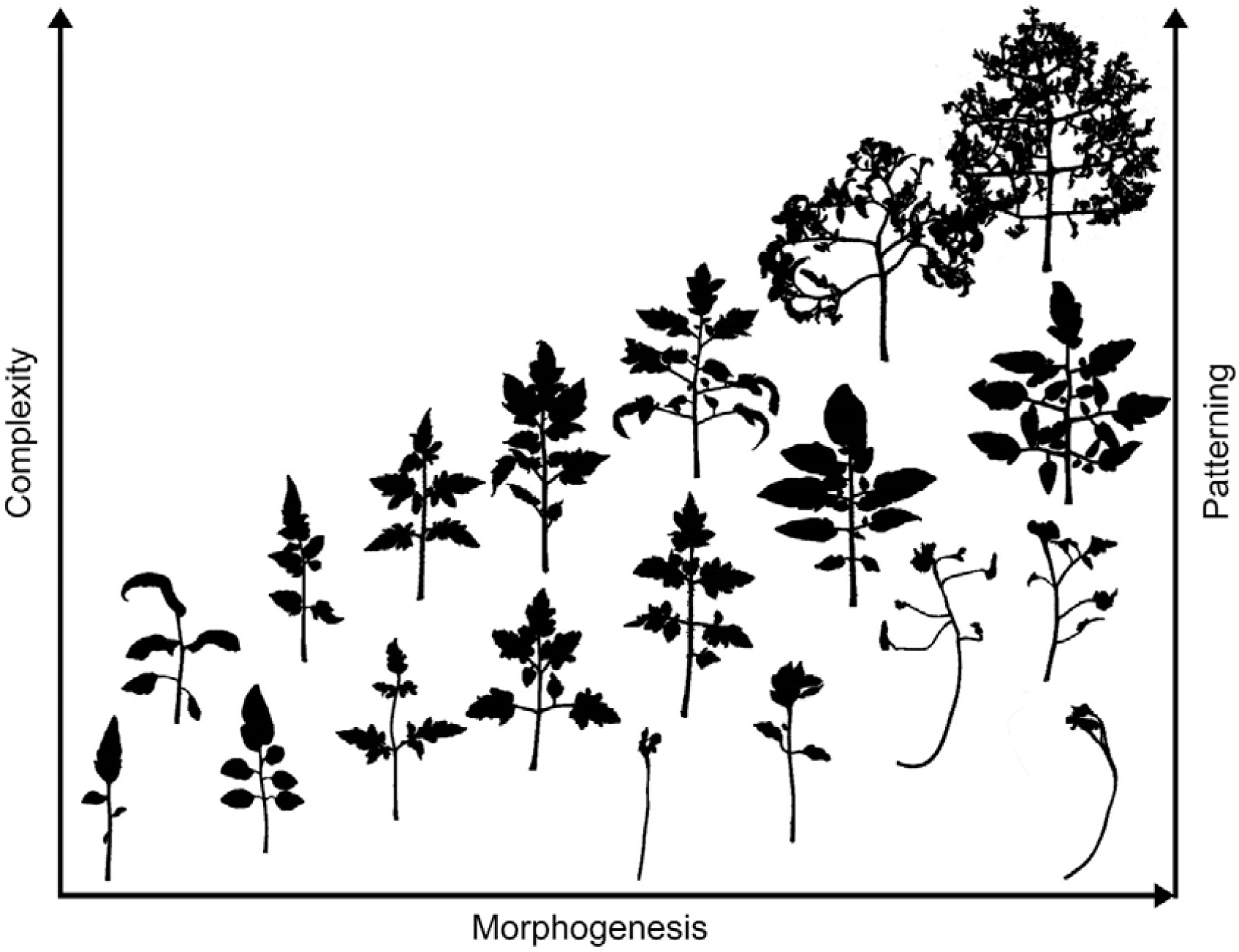
A schematic model for the proposed interaction between morphogenesis and patterning in tomato compound leaves. The diverse leaf forms shown in this study are placed in the given space according to their relative morphogenetic and patterning activities, and the resulting overall complexity. Leaf complexity is defined by the total number of leaflets and the degree of lobing at the leaf margin. This complexity depends on both a prolonged window of morphogenetic activity and on coordinated patterning events. The morphogenetic activity is defined by regulators such as: LA, CLAU, KNOXI, FALS and TF. The longer the morphogenetic activity, the more leaflets and lobes formed (Top right). Coordinated patterning is mediated by growth promoting factors such as SlMP and LYR and boundary specification factors such as GOB. Their activity is limited to the transient morphogenetic window. Reducing growth, boundaries or both, leads to a reduced leaf complexity. Interestingly, the morphogenetic activity affects leaf lobing less than it affects overall leaflet number, suggesting that these two patterning events follow at least partially different developmental programs. The diversity in tomato leaf shape is derived from tuning morphogenesis and patterning. Leaf size also depends on these two components and is partially correlated with leaf complexity.

### Multiple regulators of morphogenesis and patterning

Leaf development is a continuous process composed of several partially overlapping stages (Dengler & Tsukaya, 2001; Hagemann & Gleissberg, 1996; Poethig, 1997;). Shortly after leaf initiation from the periphery of the SAM, the primordium margins undergo primary morphogenesis, during which the main patterning events, including the initiation of marginal structures, take place. The initiation of separate blade units depends on the specification of three differential growth domains: the growth, intercalary and boundary domains (Berger *et al*., 2009; Bilsborough *et al*., 2011; Hagemann & Gleissberg, 1996; Israeli *et al*., 2019; Poethig, 1997). Genetic evidences suggest that specification and maintenance of these domains is regulated by the activity of several, partially overlapping regulators. For example, three main boundary specification regulators were identified in tomato: GOB/CUC2, LATERAL SUPRESSOR (LS) and POTATO LEAF (C). While GOB and LS were shown to act in the same genetic pathway (Rossmann *et al*., 2015), GOB/CUC2 and C likely act via different pathways to specify the boundary domain, as *gob c* double mutants enhance the single mutants and have a completely entire margin with no formation of leaflets (Busch *et al*., 2011). We have previously shown that several A-ARFs have partially overlapping function in promoting growth from the leaf margin (Israeli *et al*., 2019). Here, we show that SlMP and LYR act in parallel pathways to promote the growth domain. While *SlMP* is expressed throughout the leaf margins (Israeli *et al*., 2019), *LYR* is more specifically expressed at the sites of leaflet initiation (David-Schwartz *et al*., 2009). In addition, *LYR* expression is not affected in the *slmp* mutant (Israeli *et al*., 2019). Therefore, LYR is probably not regulated by SlMP but acts downstream of a different auxin mediator and/or by another, yet unknown factor. Therefore, several independent factors regulate the specification of each domain, and their activities only partially overlap, such that in the absence of one of them, patterning is compromised, but the basic structure is retained. This partial overlap thus contributes to the balancing between robustness and diversity (Abley *et al*., 2016; Israeli *et al*., 2019).

### Conservation and divergence in leaf patterning

Several lines of evidence suggest that NAM/CUC and class I KNOX (KNOXI) proteins positively affect the expression and activity of each other. In Arabidopsis, tomato and Cardamine KNOXI proteins act downstream of CUC in the establishment and maintenance of the SAM. *KNOXI* expression is activated by CUC and is absent in *cuc* mutants, which lack a SAM (Aida *et al*., 1999; Brand *et al*., 2007; Blein *et al*., 2008; Hay *et al*., 2006; Rast-somssich *et al*., 2015). Here, we find that compromised GOB/CUC2 activity substantially suppresses the increase in leaf complexity caused by *KNOXI* overexpression. Similarly, in Cardamine leaf development, elevated KNOXI activity leads to increased leaf complexity (Hay & Tsiantis, 2006), and reduced CUC activity suppresses this effect of KNOXI (Blein *et al*., 2008). This suggests that the relationship between CUC and KNOXI is conserved in several species and developmental processes (Alvarez *et al*., 2016; Floyd & Bowman, 2010). Interestingly, Arabidopsis CUC2 and CUC3 were dispensable for marginal elaboration, unlike tomato (Alvarez *et al*., 2016). Therefore, the regulation of marginal elaboration is distinct between tomato and Arabidopsis. However, other factors appear to mediate marginal elaboration in both Arabidopsis and tomato. Mutations in genes encoding growth-promoting factors such as WUSCHEL-related homeobox (WOX1) and (PRESSED FLOWER) PRS suppressed the indeterminate margin caused by miR-TCP-NGA overexpression (Alvarez *et al*., 2016). These factors were shown to act downstream of auxin and SlMP (Guan *et al*., 2017). In tomato, *slmp* mutants are shown here to suppress the enhanced complexity caused by prolonged marginal activity in mutants such as *la-6/tcp4, clau, Me, bip* and *Pts*. Therefore, growth promoting factors have conserved roles in mediating marginal activity in both tomato and Arabidopsis. Mutations in differentiation promoting factors such as LA/TCP4 and CLAU and plants with increased activity of morphogenetic promoting factors such as KNOXI differ in their phenotypes, as well as in their interactions with *slmp* and *gob*. *slmp* and *gob* reduced leaf complexity of *la-6* and *clau* to a similar extent as they did in the wild type. Conversely, *slmp* and *gob* phenotypes were partially enhanced by *Me/*+ and *Kn1* overexpression, suggesting that these factors interact in additional manners.

### Context-dependent interaction

A gradient of TCP expression correlates with leaf maturation, and leaves with reduced TCP activity produce larger and more compound leaves, while leaves with increased TCP activity are smaller and simpler (Alvarez *et al*., 2016; Efroni *et al*., 2008; Koyama *et al*., 2010, 2017; Nikolov *et al*., 2019; Ori *et al*., 2007; Palatnik *et al*., 2003). While TCP3 negatively regulates CUC expression (Koyama *et al*., 2007), loss of CUC activity can only partially explain the reduced leaf complexity of the dominant and smaller *La-2* gain-of-function mutant, because *gob-3* leaves are comparable in size to those of the wild type (Berger *et al*., 2009; Brand *et al*., 2007). The relationship between TCP and CUC is also connected to the age-dependent change in leaf shape, termed heteroblasty (Kerstetter & Poethig, 1998). Both simple and compound leaves display heteroblasty (Naz *et al*., 2013; Rubio-Somoza *et al*., 2014; Shleizer-Burko *et al*., 2011;). In tomato and other Solanum species, *LA/TCP4* expression increases earlier in early leaves relative to later leaves, correlating with a gradual increase in leaf complexity in later leaves (Shleizer-Burko *et al*., 2011). In Arabidopsis and Cardamine, higher *TCP* expression in early leaves delimits the activity of CUC via MIR164 regulated and non-regulated pathways (Rubio-Somoza *et al*., 2014). With plant maturation, the inhibitory effect of TCP on CUC activity is reduced, releasing CUC to increase leaf complexity. It will be interesting to examine whether a similar age-dependent interaction exists between LA/TCP4 and GOB/CUC2 in tomato. The negative interaction between LA/TCP4 and GOB/CUC2 may underlie the retained effect of GOB/CUC2 and MIR319 overexpression on leaf lobing in backgrounds with a short morphogenetic window. The age-related changes in *SlMP* expression and phenotype shown here suggest that the window of SlMP activity changes in successive leaves, likely responding to the changing dynamics of TCP activity.

In addition, GOB/CUC2 and auxin have distinct effects on leaflet and lobe patterning, and their interaction with the morphogenetic window also differs between these contexts. While the effect on lobing appears similar to Arabidopsis, this is not the case for leaflet patterning. Therefore, common tools are combined differently to pattern leaflets and lobes. This is consistent with the view that leaflets are more similar to simple leaves than the entire compound leaf (Bharathan *et al*., 2002; Efroni *et al*., 2010; Poethig & Sussex, 1985; Runions *et al*., 2017).

### Tweaking agricultural traits

Phenotypic diversity has been of great interest in research and agriculture for many years (Eshed & Lippman, 2019; Theophrastus, 1916). Domestication and breeding themes have selected genetic variants with beneficial traits such as a determinate growth habit, early flowering, fruit size, non-shattering seed dispersal and non-dormant seeds (Abbo *et al*., 2014). Many of these traits are related to maturation and differentiation rate, and are controlled by plant hormones, such as Florigen and Gibberellin (Boden *et al*., 2015; Cong *et al*., 2008; Eshed & Lippman, 2019; Lemmon *et al*., 2018; Müller *et al*., 2015; Pourkheirandish *et al*., 2015; Soyk *et al*., 2019; Studer *et al*., 2011; Zhu *et al*., 2013). The current results highlight the potential of tweaking growth patterning within the context of a maturation program to achieve shape diversity. This may inform future studies in other agriculturally important developmental processes that are regulated by patterning and maturation, such as inflorescence structure, flower and fruit development (Eshed & Lippman, 2019; Park *et al*., 2012, 2014; Rodríguez-Leal *et al*., 2017;). In addition, future breeding programs may use this approach to increase yield in a range of crops by combining subtle changes in relevant developmental traits.

## Acknowledgements

We thank Aya Refael Cohen for initial analysis of the *La-2* genetic screen and members of the Ori group for continuous discussion and support.

## Author contribution

A.I, N.O., and O.B.H. design of the research; A.I., O.B.H., Y.B., I.S., H.B.G., S.H.S., M.B., I.E. and N.O. performed the experiments, collected, analyzed and interpreted the data; A.I. and N.O. wrote the manuscript with input from all authors.

